# Deciphering the histone code to build the genome structure

**DOI:** 10.1101/217190

**Authors:** Kirti Prakash, David Fournier

## Abstract

Histones are punctuated with small chemical modifications that alter their interaction with DNA. One attractive hypothesis stipulates that certain combinations of these histone modifications may function, alone or together, as a part of a predictive histone code to provide ground rules for chromatin folding. We consider four features that relate histone modifications to chromatin folding: charge neutralisation, molecular specificity, robustness and evolvability. Next, we present evidence for the association among different histone modifications at various levels of chromatin organisation and show how these relationships relate to function such as transcription, replication and cell division. Finally, we propose a model where the histone code can set critical checkpoints for chromatin to fold reversibly between different orders of the organisation in response to a biological stimulus.

## Introduction

The genetic information within chromosomes of eukaryotes is packaged into chromatin, a long and folded polymer of double-stranded DNA, histones and other structural and non-structural proteins. The repeating units of the polymer, the nucleosomes, are 147 base-pairs (1.75 turn) of DNA wrapped around an octamer of 4 histone proteins [1, 2]. Nucleosomes are thought to be further compacted into a higher order 30 nm chromatin fibre by linker histone H1 [3]. The structure of nucleosomes can be altered post-translationally by the small chemical modifications of histone protein [4, 5]. Subsequently, one can characterise the organisation of chromatin into three interrelated categories: (1) the basic building blocks, (2) the functional structure of chromatin and (3) the higher order spatial arrangement of chromatin.

The two classical building blocks (Figure 1A, first column): beads-on-a-string and 30 nm chromatin fibre have been extensively studied previously [1, 6–8]. Regarding the intermediary level of compaction, chromatin can display several configurations (active, repressed, inactive) depending upon enrichment of a particular histone marks (Figure 1A, second column). At a higher order (Figure 1A, third column), chromatin can be either described as a bimodal heterochromatin/euchromatin model (condensed and open regions, respectively), chromosome territories [9] or as a very condensed structure in the case of the metaphase chromosome. Chromatin fibres indeed present a variety of sizes [10] and of shapes ([11] shows a solenoid model of chromatin), while recent studies attempt to challenge their existence [12], hinting that the hierarchy beads-on-a-string/fibres/domains might be much more complicated and diverse than we currently think.

**Fig. 1.**
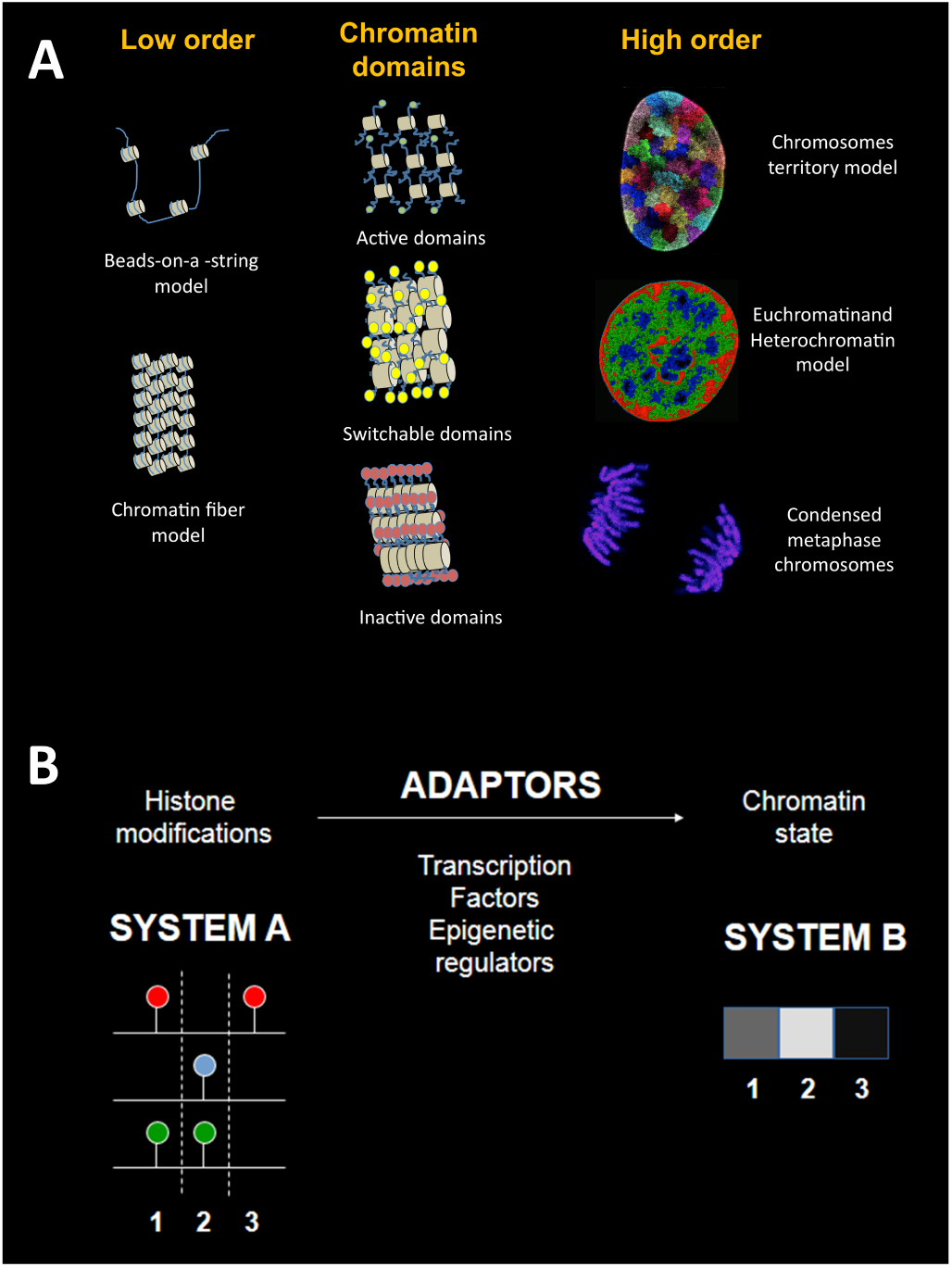
Chromatin spatial organization and relationship with the histone code. (A) The spatial organization of chromatin can be studied at three levels: at the lowest orders, which include the beads-on-a-string model and the chromatin fibre (left column), at the level of functional chromatin domains (middle column) and high order chromatin patterns (right column). The organization of chromatin domains can be modelled using various post-translational histone modifications. Depending upon the kind of histone modification, a nucleosome is enriched with, chromatin can be either highly condensed, in an open conformation or switch between these two extreme forms. The higher order chromatin patterns can either be viewed from: the chromosome territory model of DNA organization, the over-simplified bimodal classification of condensed and open chromatin as hetero- and euchromatin respectively, and the highly condensed configuration of metaphase chromosomes. (B) The components of the histone code. On the left, the input system A, made of combinations of histone modifications (coloured pins). One unique combination can be found at three different positions 1,2,3, in the genome. The combinations of the input system A is translated into components of the output system B, with a black and white colour-scale. The translation happens via adaptors such as transcription factors or epigenetic regulators that recognise specific histone modifications to change chromatin state.

On the functional side, it has been shown that biochemical changes made to specific histones tails are associated with different condensation levels of chromatin. For instance, trimethylaton of lysine 9 on histone 3 (H3K9me3) is usually associated with condensed chromatin and centromeric regions [15, 16]. On the other hand, trimethylaton of lysine 4 on histone 3 (H3K4me3) is strongly enriched at promoter regions of active genes where chromatin exists in an open conformation [17, 18]. Chromatin can also exist in a switchable state between these two extreme forms if histone H3 is trimethylated at lysine 27 (H3K27me3) [19]. H3K27me3 is a histone mark characteristic of repressed genes.

The number of ways histone proteins can be modified is quite extensive and can explain the vast majority of possible combinations that can lead to various functional outcomes. The relationship between these combinations and the function they perform is referred to as the "histone code" [20–22]. Previously, biological codes have been thoroughly described [23–25]; they comprise of an input system, made of signs with no biological function in itself that are translated into organic output functions via adaptor molecules. In the example of the genetic code, the inputs are codons, the adaptor is the translation machinery and the outputs are amino acids. In the case of the histone code, combinations of histone modifications at a given position on the genome constitute the input system, the adaptors are the epigenetic regulators (for instance enzymatic complex Ezh2) that bind the modifications and outputs are chromatin features such as the level of chromatin compaction or gene expression (Figure 1B).

#### Outlook

*The urge to discover secrets is deeply ingrained in human nature; even the least curious mind is roused by the promise of sharing knowledge withheld from others. Most of us are driven to sublimate this urge by the solving of artificial puzzles devised for our entertainment. Detective stories or crossword puzzles cater for the majority; the solution of secret codes may be the hobby of the few.*

J. Chadwick, The Decipherment of Linear B [13].

Citation inspired from The Code Book by Simon Singh [14].

From the day/night dichotomy to the genetic code, nature is full of symmetric, antagonistic exemplars and patterns. One such example is the organisation of structurally distinct chromatin states (active, inactive) on a single chromosome. In this article, we try to show how simple combinations of essential elements such as histone modifications can participate in sophisticated cellular features such as the structure of the genome. Here a code is identified, where an input system (histone modifications) is translated into an output system (chromatin states) via adaptors (epigenetic regulators or transcription factors). Such a code has a distinct importance in gene regulation and consequently for the cellular phenotype.

Here, we present supportive evidence for a combinatorial occurrence of histone modifications and its consequence for higher-order chromatin folding. We start with a brief historical introduction to chromatin biology. We then show the correlation among histone modifications at the level of nucleosomes, regulatory regions, TADs and chromosomes. Finally, we propose a model showing that these distinct chromatin domains can co-exist on a single chromosome.

## Histone modifications, a large vocabulary of words

Histones were first identified as a fundamental component of the nucleus in the early days of molecular biology [26, 27] and were described as circular structures responsible for compacting DNA [1, 6, 28]. They were soon associated with DNA periodicity and compaction [6, 29–32], where DNA coils around an octamer of four core histone proteins (H2A/H2B dimer and H3/H4 tetramer, Figure 2A) [1, 33]. The histone core with DNA was termed as nucleosome and is known to be the first level of chromatin compaction. Early experiments showed that histones, via their N-terminal tails, can experience modifications such as acetylation and methylation, that can further alter the compaction of chromatin [4]. The nomenclature of histone modification were named the following way: in the case of H3K9me3, H3 refers to the core histone protein, K refers to the amino acid, the number 9 indicates the position of lysine residue from N-terminal end of the amino acid tail of histone protein and me3 refers to the type of modification on the lysine residue (Figure 2B).

**Fig. 2.**
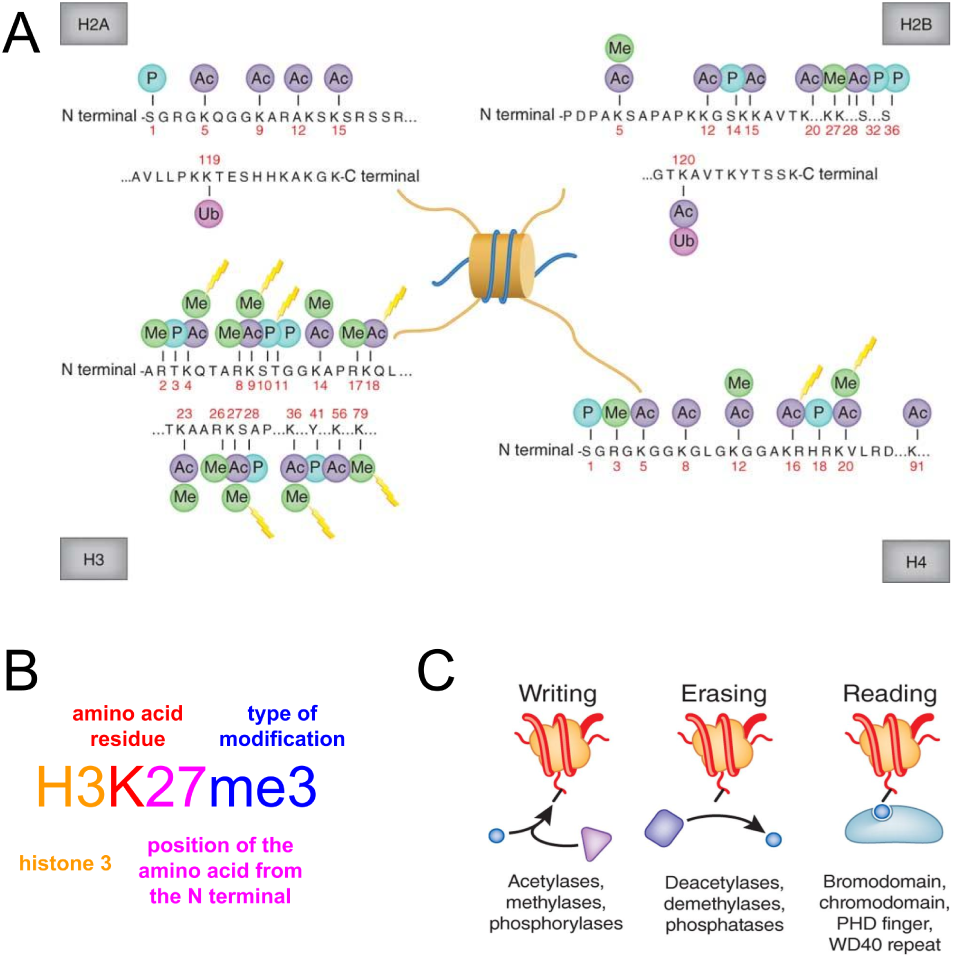
Histone modifications are major biochemical features of chromatin. Histone can experience various post-translational modifications in their protruding N-terminal tails but also within the C-terminal regions. These changes affect not only the overall compaction of chromatin but also gene expression. (A) The principal modifications on the four core histones: H2A, H2B, H3, H4. Two patterns worth noting here is the arrangement of lysines (K4, K9, K14, K18, K23, K27) and occurrence of Lysine (K), Serine (S) and Arginine (A) at (9, 10, 11) and (27, 28, 29) position on the amino acid tail of histone H3. Modified from [40] with permission. (B) Nomenclature of histones post-transcriptional modifications. The different features are the type of histone (H3 in our example), the amino acid and its position (Lysine at position 27) and the chemical modification, here a trimethylation, or triple methylation of the lysine 27 from the N-terminal end of the amino acide tail. (C) Different enzymes lead to writing, erasing and reading of chemical groups on the tails of histone proteins. Writers are enzymes that introduce various chemical modifications to histones. There are about 100 different enzymes that can write acetyl, methyl and phosphoryl groups (circles) to individual amino acid residues. Erasers are the enzymes that remove histone modifications and perform the opposite task of writers. Readers are enzymes that recognise specific chemical groups on the amino acid residues. For instance, bromodomains engage with acetylated lysines (modified from [41] with permission). Ac, acetylation; Me, methylation; P, phosphorylation; Ub, ubiquitination.

Over the course of the last decade, amino acids have been shown to experience several modifications, of at least twelve types: acetylation (lysine), methylation (lysine and arginine), phosphorylation (serine and threonine), sumoylation (lysine), ubiquitylation (lysine), ADP ribosylation, butyrylation, citrullination, crotonylation, formylation, proline isomerization, propionylation [34–39].

At a fundamental level, most of these modifications are neutralizing the net charge between DNA and histone proteins, and much of the gene regulation is a by product of effective net charge in the environment [44, 45]. Hence a combination of marks must work in synergy to provide sufficient proof reading so that no gene is accidentally turned on or off [46– 48]. Lastly, the combination of histone modifications are stably maintained across many species and protein motifs have evolved to recognise histone modifications [49].

We consider four measures which characterize the function of histone modifications

1. charge neutralisation
2. molecular specificity
3. robustness
4. evolvability

### Charge neutralisation

Histone modifications are usually modified via enzymes that can neutralise the excess of charge on the DNA [50, 51]. A nucleosome core particle has an overall residual charge of *−*150*e* (DNA contributes *−*294*e*, and histones contribute +144*e*) and is therefore electrically not neutral [52]. Tails can attract negatively remote DNA molecules (the negatively charged due to phosphate oxide groups) to induce regional DNA compaction. Subsequently, the folding of DNA is also highly dependent on the positive counterions in the environment [53, 54].

Early experiments showed that acetylation (CH3-CH2-) or methylation (CH3-) of nucleosomes occur respectively in 50-60% and 40% of histones and can lead to increase and decrease of polymerase activity, respectively [44]. Methyl groups are produced during the metabolism of methionine, a subpart of the B12 vitamin circuit while acetylation is generated from the acetyl-Coenzyme A [55, 56]. Histone acetylation impairs the affinity of histones to DNA via charge antagonism, reduction of compaction and subsequent recruitment of certain factors such as SIR at telomeres or TAF1 at promoters, inducing activation of transcription [57–59]. This property is accompanied by the higher mobility of histones along DNA [60]. Differently, methylation is associated to compaction of chromatin and so to reduced transcription, with reduced histone mobility [44, 60, 61].

### Molecular specificity

The histone modification sequence is orchestrated by the enzymes that either deposit, read or remove the marks, and serve as the adaptors molecules essential to biological codes [23]. Enzymes that deposit marks are diverse and reflect the large spectrum of modifications available in nature (Figure 2C). Well-known examples are of histone acetyl-transferases (HAT) which transfer the acetyl group from acetyl-Coenzyme A to the tail of histones, and histone deacetylases (HDACs) which remove the acetyl groups, most notably upon chromatin compaction. Other complexes involved in histone regulation include the Trithorax complex (Trx) which is involved in methylation of histones and Polycomb (PcG) with module PRC2 involved in the deposition of H3K27me3 mark via methyltransferase Ezh2 [62, 63].

Additionally, the H3K27me3 mark is bound by the PRC1 complex that can mono-ubiquitinate histone H2A on lysine 119 (H2AK119Ub1). This process is known to silence genes and is an excellent example of a combination of marks found at a genomic site with an impact on the function. In contrast, transcription factors with bromodomains are known to bind acetylated lysines [64] and induce gene activation via acetyl-transferase activity [65].

### Robustness

Robustness is the capacity of living systems to preserve a given function during evolutionary times. Many important core features of organisms are robust and found in a wide variety of species; for instance, the skeleton is found in all vertebrates and can be considered a robust feature. Small or big structural variations are present in all groups, but the template remains the same. Robustness seems to imply two features: cooperation and redundancy; cooperation of the different basic molecular elements of the system, including quality control checkpoints [66] and a high redundancy of the molecular elements, in order to resist to small changes. In the example of the skeleton, many different genes are involved, each of them participating to build the scaffold (cooperation). Nevertheless, many genes perform the same function so if one gene fails or fluctuates, several other ones can take over to perform the same function (redundancy).

Overall, robustness is a requirement for all fundamental systems of life. First, all the core biological codes such as the genetic code, which can produce proteins despite occasional changes [67]. Similarly, the histone code seems to feature properties favouring robustness. As we will see later, very similar histone modifications such as acetylations or methylations act in concert to form the histone code (cooperation), which potentially means that the loss of one mark does not necessarily affect the outcome output of the code very strongly (redundancy). For instance, for a given genomic region, active promoters, different combinations are possible, with small change (Figure 4). Such a redundancy stems in the high similarity of nucleosome structure across kingdoms Archaea and Eukaryotes [68] and the fact that most well-known histone modifications are found across Eukaryotes, including human, mouse, fly and C.elegans [69]. For instance, H3K9me3 is found to be present in vertebrates, invertebrates and yeast [70, 71]. Functions of enzymes associated with histone modifications re-modelling seem to be very conserved across Eukaryotes [72, 73]. Subsequently, the histone code provides a robust system that is not prone to mistakes and delivers a reliable answer to stimuli, therefore, seems to fill all requirements of a robust system.

### Evolvability

A consequence of robustness is that small variations are usually harmless, a property that opens doors to small changes through evolution, with various consequences. Evolvability of the histone code indeed does not operate at the level of histone sequence itself, as histone sequences are found to be very conserved. Nevertheless, minute changes in histone associations are found between relatively close species such as chimpanzee and human, or primates and rodents [69, 74]. Other marks such as H3K4me3 seem to have different patterns in complex organisms compared to rather simple distribution in yeast.

As a result of these different features, the histone code can be viewed as a stable system (notion of robustness) with enough flexibility that provides a basic ground to generate new functions by accumulation of slight changes of its elements (notion of evolvability).

## Histone code dictates combinatorial associations between histone modifications at the nucleosome level

Many histone modifications have been proven to function in a concerted manner with other marks. Theory by Jenuwein and Allis hypothesizes that these combinations serve as elementary signs which are translated into instructions by chromatin protein complexes to regulate the genome [21]. Genomic approaches such as ChIP-Seq have identified the precise localization of proteins on DNA, and even more strikingly, have been able to distinguish between different modifications due to highly specific antibodies against mono- di- or trimethylation, among others (see [75], a pioneering example). The findings in these studies show that histone modifications of different classes are found simultaneously at the same genomic positions, id est in the same nucleosome. This is the result of an amplification process, for example, H4K20me1 can be recognized by enzyme Suv4-20h1 which triggers a second methylation event on the same lysine, generating a H4K20me2 mark [76, 77]. Another example is the PcG complex involved in gene silencing, which is made of the association of complexes PRC1 and PRC2. PRC2 is specialized in depositing the H3K27me3 mark [62, 63], while PRC1 recognizes H3K27me3 [78] and subsequently ubiquitinates lysine 119 of histone H2A via protein Ring 1B E3 ubiquitin ligase [79], amplifying the signal.

More recent studies have demonstrated that variants of the PRC1 complex can come in first to deposit H2A ubiquitination, which is thereafter recognised by PRC2 to finally deposit H3K27me3 [80, 81]. Histone modifications do not need to be exactly on the same histone protein to act synergistically or on neighbor nucleosomes, providing that the different modifications acting synergistically are close together in 3D space [82]. The combinations of histone marks affect transcription and so indirectly cellular events which are important for cell and organism identity. For instance, during interphase, phosphorylation at H3S10 associates with H3K9 and H3K14 acetylation and induces chromatin relaxation [83]. During mitosis, DNA needs to be packed and H3S10 phosphorylation heavily triggers compaction. Two combinations with the same mark participate to two different cellular outcomes.

The amount of information carried by the overall collection of modifications can be probed by checking the correlation of marks between individual histones, using data from ChIP-Seq [42]. The data reveals a global redundancy factor of about 1:7 with five different clusters using 39 different marks (Figure 3A). As a result, the number of possible functions that can be directed by the system, even if the number of marks is high, is relatively low, which is confirmed by genomic studies [84]. An insightful model has also shown that a simple two-marks histone code can already provide a wide panel of function, hinting that few core elements can lead to a large combination of genomic functions [85, 86].

**Fig. 3.**
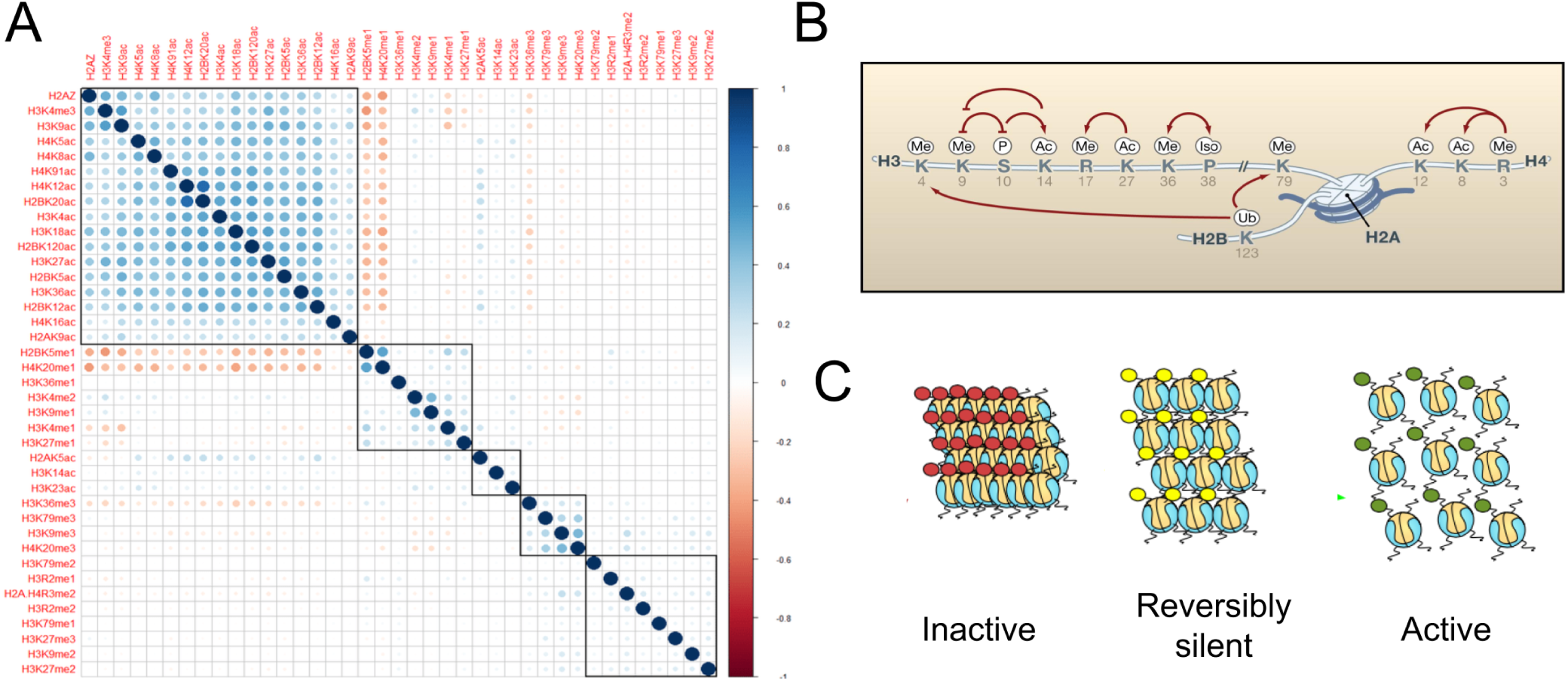
A binary combinatorial histone code at the level of nucleosomes. (A). Different clusters of histone modifications is revealed by hierarchical clustering. Correlation between modifications are displayed with a color scheme, red to blue from negative to positive correlation coefficient. Clusters are identified by black boxes (figure from [42], courtesy Kernel Press, Mainz). (B) Different cis and trans patterns of histone crosstalk (figure from [43], reprinted with permission). (C) The impact of three primary histone modifications on chromatin compaction. H3K4me3-rich nucleosomes are associated with active chromatin in an open conformation, H3K27me3-rich nucleosomes are associated with repressed chromatin, while H3K9me3-rich nucleosomes are associated with highly-compacted, inactive chromatin.

Direct experimental evidence for associations of histone modifications was brought by studies using co-marking of histone modifications [87] and genomic experiments associated to microscopy (co-occurring iChIP in [88]). The single-nucleosome experiments confirm very well-known histone code rule to happen on isolated nucleosomes, such as the bivalent state (H3K27me3/H3K4me3), which is typical of embryonic stem cells. An example of a cis and trans crosstalk within a nucleosome is shown in Figure 3B.

Advance microscopy techniques [89–92] have been powerful at seeing the distribution of histone modifications in the nucleus [93–95]. In particular, single molecule localisation microscopy can discriminate between regions painted with H3K4me3 and H3K27me3 [96, 97]. These marks are highly associated with the general compaction level of chromatin (Figure 3C). Overall, one usually distinguishes between euchromatin and heterochromatin, the first being opened, in an "active" state prompt for genes to be transcribed, while the latter is in a repressed state [98–100].

## Histone code associations with genome function and structure

Historically, the association of histone modifications with transcription is known since long [44, 103]. Regarding modification associations, studies have revealed that an increase in the complexity of the histone make-up results in progressive changes in the gene expression [104], with acetylation to be globally associated to increase in transcription and deacetylation with decrease in transcription. To our knowledge, H3K36 methylation by the SET2 complex (co-transcriptionally binding to elongating Pol II) targets Rpd3S deacetylases to suppress cryptic transcription start sites within genes [105, 106].

ChIP-seq experiments have helped to draw a more refined picture of the relationships between histone modifications and gene function [101]. Hidden Markov models were employed to guess where a histone modification combination (or “chromatin state”) starts and ends. After testing many combinations of parameters, the model was found to compromise 51 states, which is a surprisingly large number. Most of these states are found to be associated with specific genomic regions. 11 out of the 51 states fall in promoter regions, which is a significant enrichment when considering the low amount of promoter regions in the genome compared to non-coding regions (Figure 4A). Promoter regions are associated with acetylated marks, all of which show a similar bimodal profile around transcription start sites (TSSs) (Figure 4B). Transcribed regions are associated with particular kinds of methylation, entirely different from the ones observed in repeated regions, namely methylations at lysine 4 of histone 3 (H3K4me1 and H3K4me2) along with methylation on lysine 79 of histone 3. Finally, the table shows that repeated regions display a unique combination of acetylation and methylation which is quite predictable. The rules illustrated in Figure 3A have since been extensively described in a broad range of tissues studied in the large consortium devoted to epigenetic studies called ENCODE [71].

**Fig. 4.**
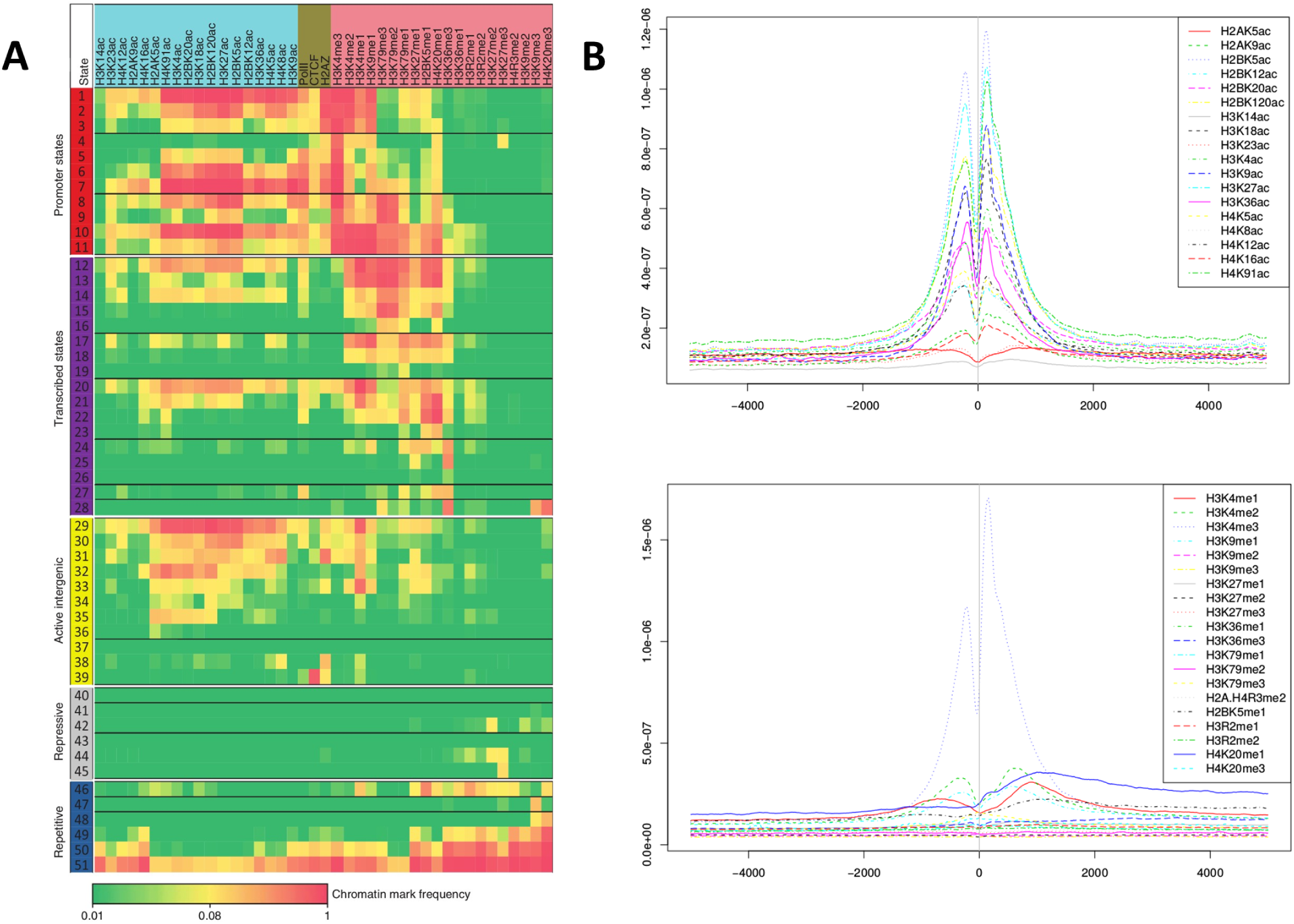
The histone code at the level of genomic features. (A) The combinations of histone modifications and their associations to various genomic states (figure modified from [101], reprinted with permission). Here, the genome is partitioned into five broad regions: promoter states, transcribed states, active intergenic, repressive and repetitive state. A combination of histone marks is associated with each genomic state. The relative degree of association is indicated by a colour scale spanning frequency values between 0 and 1. (B) Profile of histone modifications around the transcription start site (TSS). On top panel, profile for acetylation marks; below panel: methylations. X-axes: Distance to TSS. Side with negative values are upstream from the gene, side with positive values downstream, irrespective of the strandness of the gene. Y-axes: Normalized read counts. Profiles were generated by us with a R script [102].

Histone modification is indeed only meaningful in synergy with other genomic features, for instance, methylation of cytosines at CpG sites. DNA methylation is known to be associated with gene silencing. For this to happen, histone methyltransferase HP1 recognises methylation events on lysine 9 of histone 3. During DNA replication and after binding, HP1 recruits the DNA methyltransferase DNMT1 that induces hemi-methylation of neighbour cytosines, inducing gene promoter inactivation [108]. Differently, DNA methylation is known to stimulate the deposition of repressive mark H3K9me3 and induce DNA compaction at promoters [109], via binding of methylated cytosine by MeCP2 and deposition of the mark by coupled enzyme Suv39h1 [110].

The impact of histone modifications on transcription and other cellular functions is largely based on the influence of chromatin structure. At a nanoscale, methylation or deacetylation of histone tails ensure the stability of nucleosomes [50]. At a gene scale, the makeup is thought to be locally uniform, guiding folding properties, for instance in the region of Hox genes [67]. Areas with similar epigenetic make-up may fold together while repelling other parts of different make-up, thus contributing to the formation of locally coherent topological chromatin domains in the size range of 0.5-1 Mb, or TADs described by recent chromatin conformation capturing approaches [84, 111]. Indeed, we have recently shown that histone modifications cluster at genome scale using localisation microscopy [112]. Previously, it was shown with the confocal microscope, that H3K4me3-rich, H3K27me3-rich and H3K9me3-rich regions partially segregate (for instance in [113]). On our localisation images, interphase and M-phase chromatin could also be roughly divided into H3K4me3-rich, H3K27me3-rich and H3K9me3-rich regions of highly different patterns. These marks are known to be respectively associated with active genes, inactive genes and deeply repressed chromatin [114].

Along with microscopy, genomics has been the most powerful at characterising the relationship between modifications of histones and structure. The HiC method has helped to identify regions of structurally complex DNA folding, whether loose or compacted, that are thought to be functional units [111]. ChIP-seq of histone modifications has shown that chromatin domains described with the HiC method can be associated with combinations of histones (Figure 5A). In mammals, domains related to active chromatin usually show H3K4me3 and acetylated histones H3 and H4 at the promoters of genes, while inactive domains display genes silenced and mark H3K27me3. Differently, H3K27me3-rich domains have the potential to be activated on demand, depending on the tissue, and so are more open than H3K9me-rich domains [115]. Finally, domains associated to H3K9me2/H3K9me3 are very compact and constitute a significant portion of chromatin that is never active and is structurally very condensed. Figure 5B shows a localisation microscopy picture, the patches of active, inactive and repressed chromatin, painted with H3K14ac, H3K27me3 and H3K9me3 [10]. From these different findings, we propose a model for chromatin domain formation (Figure 5C).

**Fig. 5.**
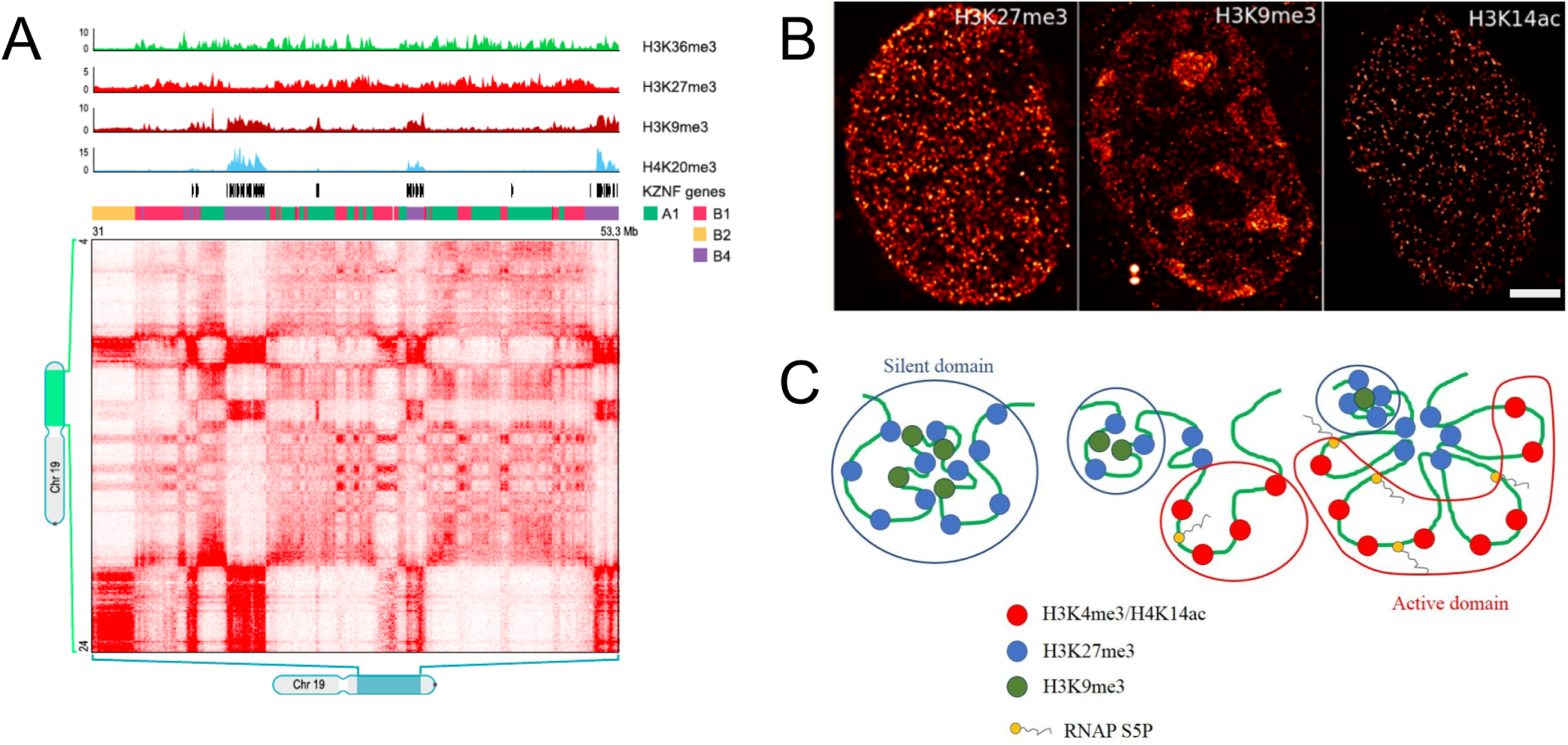
The histone code predicts the degree of folding of chromatin domains. (A) A section of chromosome 19 shows different subcompartments based on histone modifications. Here, chromatin domains with high contact frequencies correlate with histone marks associated with inactive chromatin such as H3K9me3 and H4K20me3. Such relation between H3K9me3 and H4K20me3 was also found at the nucleosomes level based on the entirely different method and cell line. This shows that relationship between histone modifications holds at various length scales. Figure reprinted from [84], with permission. (B) Imaging of histone modifications further reveals the distribution of domains associated to different histone modifications. The different histone modifications have distinct molecular signatures, probably due to different folding rules and roles in gene regulatory mechanisms. While H3K9me3 seems to be mostly enriched at chromocenters as expected, H3K27me3 is distributed more in speckle-like patterns. The acetylation patterns (H3K14ac) vary in density and are more diverse in distribution. Figure adapted from [107]. Scale bar: 1 *µm*. (C) A model portraying three stages of a Hox gene activation, with progressive suppression of a silent chromatin domain made up of H3K27me3- and H3K9me3-rich histones toward the installation of an active domain rich with H3K4me3. The change of domain happens literally at the same place, with inhibitory marks being progressively evicted from the region to be replaced by the active domain. Note the region of inhibitory marks remains at the bottom, to keep the two extremities of the loop in a condensed fashion and leave the loose active region in a relatively consistent open shape.

## Histone code across the cell cycle

During the cell cycle, the genome structure is heavily modified via epigenetic mechanisms. Nevertheless, activity and regulation of genes have to continue to a certain extent. Keeping the epigenetic makeup of chromatin during DNA condensation at the end of metaphase to switch to mitosis is relatively costly in terms of energy, so some optimal regulatory mechanisms have to happen to "memorise" the epigenetic status of the dividing cell. So far, we have considered cells in G1 or G0 phase, but the picture is rather different during mitosis, especially with respect to histone modifications. Immediately following replication, histone proteins start to accumulate on the newly-synthesised DNA to form nucleosomes, involving a specific variant of histone H3, H3.1. The late cell cycle phase has two kinds of epigenetic regulations [116], which we review briefly below.

Firstly, some marks are involved in the mechanism of cell cycle progression itself. Accumulated H4K20me1 mark at gene bodies is read and leads to recruitment of factors participating in the initiation of DNA synthesis at origins of replication such as L3MBTl1 and condensin II [117, 118]. This recognition mechanism leads to amplified events of methylation, and accumulation of marks H4K20me2 and H4K20me3 via methyl-transferases SUV4-20H1 and SUV4-20H2 to switch from S- to M-phase [119–121]. Another mark involved in the process is H3S10 phosphorylation (H3S10ph). Along with methylation at position H4K20, it is the only histone site which varies through the cell cycle. This simple two-mark regulation is enough to induce cascades of factor expression involved in further triggering chromatin compaction during mitosis [122].

Secondly, besides the epigenetic mechanisms that accompany replication and mitosis, another kind of epigenetic regulation is devoted to preserving the transcriptional status of genes so that activity can be restored in daughter cells at the end of mitosis. The view gets even more complex when considering symetric and asymetric division [123, 124]. The two marks which participate the most in this kind of remodelling are H3K27me3 and H3K9me3 [116], driving gene silencing and high chromatin compaction, respectively. A fundamental biological question is whether the epigenetic marks and so the histone code rules are preserved through replication, a mechanism named epigenetic inheritance [125–127]. In order replication to happen, the DNA becomes naked, and the nucleosomes need to be reassembled so chromatin keeps its functions in later stages. Many mechanisms are known, some involving a de novo “random” refolding which is independent of the DNA status before S-phase, and one mechanism dependent on the previous status, involving an epigenetic memory. In one model, all histone modifications are preserved and memorised on each DNA strand of the two double helices after replication. Differently, histone modifications can be partially transmitted to the daughter DNA molecules, but via copy mechanisms involving histone modifications, readers and writers recruited locally. In this case, all gaps can be filled to restore an epigenetic state very close to the one of the parent DNA molecule.

Another important question is the recycling of nucleosomes and histones. Are nucleosomes de-assembled and reconstructed in the newly synthesised DNA molecules or are they mostly transmitted in an intact form, with missing nucleosomes being de novo assembled? The answer is that in most cases, each nucleosome is transmitted intact to one of the two daughter strands, so the histone code is preserved [123, 128]. The equivalent position on the other strand is possibly filled with information coming from adjacent redundant nucleosomes via enzymes CAF-1 and ASF1 [129].

Despite costly at first sight, epigenetic inheritance may be relatively cheap to maintain with the recruitment of proper readers and writers in the vicinity of replication sites to stochastically recreate the “parental” epigenetic make-up in daughter strands. In the future, integration of modelling, genomic and microscopy features will be key to understanding the mechanisms of epigenetic inheritance.

## The association of histone code with cell identity and dynamics

Overall, genomic studies have shown that there is a substantial reshuffling of histone modifications during the cell cycle [84]. To show these associations, careful choice of FISH probes associated with H3K4me3/H3K27me3/H3K9me3 were found to demonstrate that chromatin domain stained with any of these marks anti-localise (Figure 6), a fact confirmed in Drosophila more recently, with overlaps [115]. The model of domains that we described previously is static, and in reality, chromatin domains experience a large change in embryonic development and differentiation processes of adult tissues (Figure 7). According to microscopy and genomic data during early stages of development, a heavy remodelling of chromatin structure happens. Most of the chromatin is relatively loose at first, and becomes more compacted in later stages, with associated silencing of many genes. In embryonic stem cells, a relatively high number of genes are expressed, including genes for maintenance of stem cell status, such as cMyc or Sox2 [130]. These genes are usually painted with H3K4me3 and H3K27ac marks at their promoters, prone to recruit transcription factors such as Pou5f1, Sox2, cMyc and Klf4 [131]. Many specialised genes are on the other hand showing poised promoters. These promoters have an ambivalent epigenetic status, comprising both active and inactive marks H3K4me3 and H3K27me3, respectively, and have the potential to either be activated or repressed during differentiation.

**Fig. 6.**
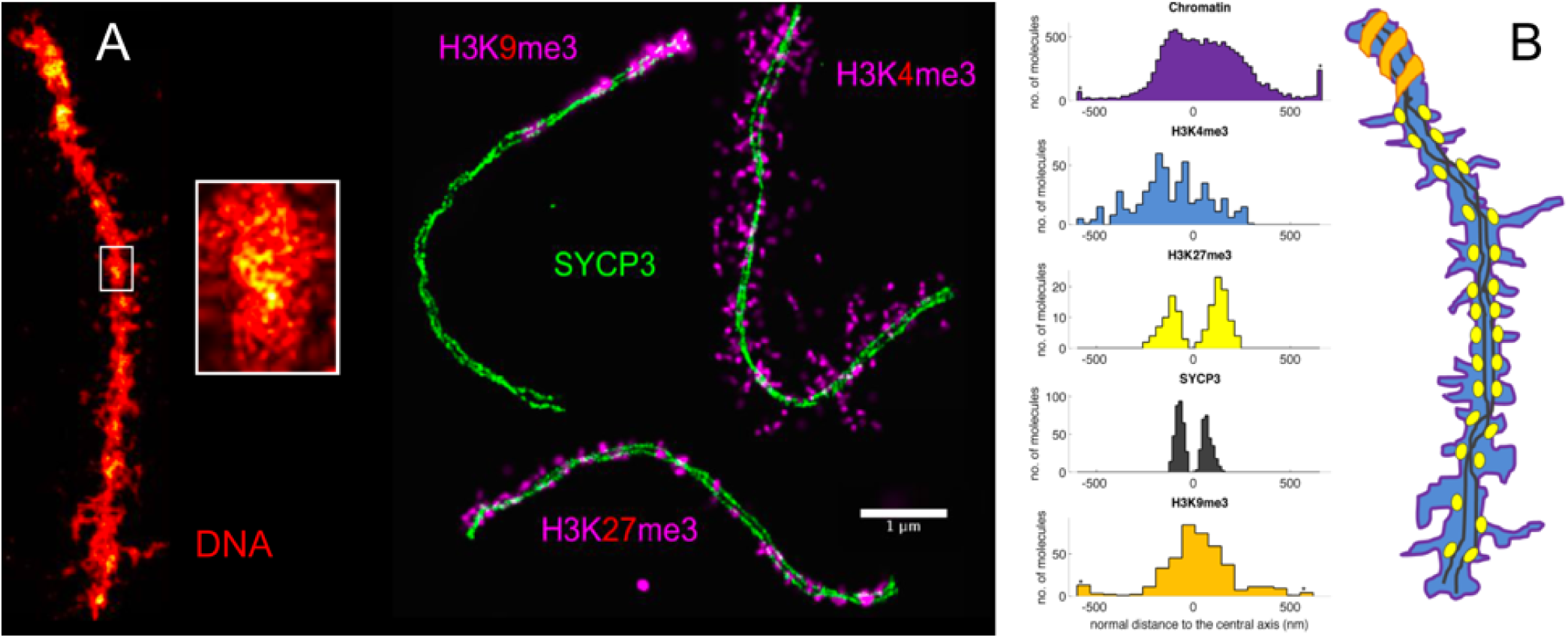
Histone code segregates functional compartments in the mieotic chromosomes. (A) Super-resolution microscopy reveals higher-order clusters of chromatin patterns along the pachytene chromosome. The DNA map of meiotic chromosomes at pachytene stage shows different condensation levels. These levels were found to be constrained by anti-correlating clusters of histone modifications (pink) along the synaptonemal complex proteins 3 (SYCP3) (green). Histone modifications associated with transcription (H3K4me3) emanates radially in loop-like structures while the histone modifications associated with repressive chromatin (H3K27me3) are confined to axial regions of the SYCP3. The histone modifications associated with centromeric chromatin (H3K9me3) are found at one end of the SYCP3 (figure and caption modified from [42, 114]). (B) A model for co-existence of different chromatin domains on a single chromosome. The two strands of the SYCP3 are displayed in black. In purple, active chromatin marked by H3K4me3 covers the entire surface of chromosomes (shown in blue). Symmetrical and periodic clusters of repressed chromatin H3K27me3 are shown with yellow spots. Inactive polar chromatin marked by H3K9me3 appears as an orange spiral pattern. The plot in the middle shows the distance distribution of DNA, active (H3K4me3) and inactive (H3K27me3, H3K9me3) regions of the genome from the central axis of SYCP3.

**Fig. 7.**
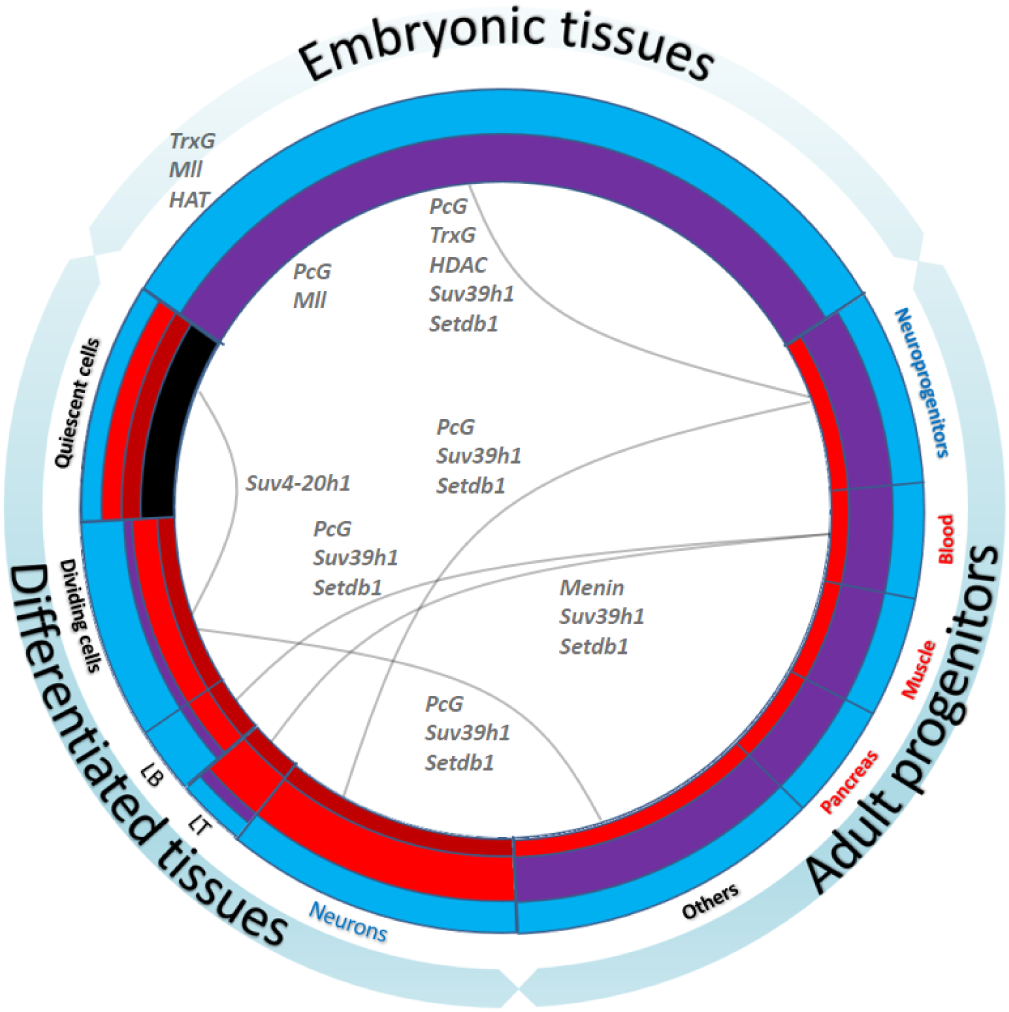
Histone code during developmental processes. The model depicts the rules of the histone code associated to major developmental and regulatory processes happening in the human body. The different colours of circle feature different kinds of histone modifications combinations with the most significant modifications are displayed. Blue: Euchromatin (H3K4me2/3 and H3K27ac, see Hawkins et al. 2010 [133]). Brown: repressed chromatin (H3K9me2/3, Ibid). Red: Inactive chromatin (H3K27me3, Ibid). Purple: Poised transcription (H3K4me3 and H3K27me3, see Gapp et al. 2014 [134]). Black: Quiescent chromatin (H4K20me2/3, see Evertts et al. 2013 and Onodera and Nakayama 2015 [132, 135]). Pathways from one genome status to another are figured by pale orange lines associated with names of chromatin modifiers or readers involved.

Another key event of cell differentiation is the increase of the proportion of densely repressed chromatin, usually associated with telomeres and nucleoli. These regions are associated with H3K9me3 and eventually H3K27me3 (Figure 6B). Cells become slowly more specialised during development, which means that their specialised metabolism focuses on expressing of only a small subset of genes with a majority of genes being put in a repressed state. A lot of genomic locations indeed become repressed, histone combination usually turns to a deacetylated state with repressed marks, such as H3K27me3. Some tissues experience bigger changes than others, for instance, T lymphocytes present an enrichment for the H3K27me3 mark compared to B lymphocytes [132].

A tight orchestration of gene regulation happens during the developmental processes associated with Hox genes, which are key in defining general body plans [136, 137]. They form clusters and are either transcribed together or sequentially during different developmental phases. In Drosophila, Hox genes are repressed at the beginning of development, under the influence from endogenous PcG complexes of the maternal source [138, 139]. With time, Hox genes are de-repressed and activated by methylation of lysine 4 of histone three while H3K27me3 is dissociated. Progression is also spatial, with Hox genes being activated from one end of the cluster to the other [140]. Furthermore, both “active” and “inactive” parts of the Hox genes cluster form separate chromatin domains (Figure 5C). Differently, in tissues with no Hox gene expression, such as brain, most of clusters are silenced and fold according to cluster positions, instead of active/inactive status, showing a deep connection between histone code, dynamics of transcriptional regulation, high-order DNA folding and consequences for body-wide phenotype.

## Conclusion

We have presented arguments for the existence of a spectrum of epigenetic features related to chromatin structure, epitomized by the term histone code. A consequence of the intrinsic redundancy of the code is robustness to small changes and therefore the message it delivers will not be very much altered if, for instance, an enzyme depositing one mark is not fully functional. Nevertheless, the position themselves where the histone modifications occur, are not prone to variations. As an example, the alteration of lysine H3K27 into a methionine is found in a certain type of glioma [141]. We can only hypothesize that similar mutations in other important modifications may have other dramatic effects for cellular phenotype and life span of organisms with either cancer or developmental defects. However, most importantly, redundancy of the code is a way to increase the probability for factors to be brought to the nucleosomes, for instance in a case of a protein complex where one protein binds to one histone modification, and another protein binds to a different histone modification. This is shown by the fact that factors from the same complex usually bind similar marks or associated marks either on the same histone or close positions. The free energy of the compound/nucleosome system will increase, increasing the probability of binding [142].

Redundancy of information provided by the histone code is an essential feature of robustness of cellular processes. As the number of histone modifications defining a given chromatin state increases, so does the probability to recruit specific factors that will change or modulate the state of chromatin. Overall, the complexity of histone modulation helps to ensure that events will happen at the right time in the right place. Though not fully understood yet, these processes are certainly a key point to understand the mechanisms that connect gene regulation and chromatin folding.

## ACKNOWLEDGMENTS

D.F. thanks the Center for Computational Sciences in Mainz for the financial support. K.P. thanks Wioleta Dudka for proofreading the manuscript.

